# Microbial virulome and resistome in deep-sea cold seeps pose minimal public health risks

**DOI:** 10.1101/2024.11.25.625267

**Authors:** Tianxueyu Zhang, Yingchun Han, Yongyi Peng, Zhaochao Deng, Wenqing Shi, Xuewei Xu, Yuehong Wu, Xiyang Dong

**Affiliations:** School of Oceanography, Shanghai Jiao Tong University, Shanghai 200030, China; Key Laboratory of Marine Ecosystem Dynamics, Second Institute of Oceanography, Ministry of Natural Resources, Hangzhou 310005, China; Key Laboratory of Marine Genetic Resources, Third Institute of Oceanography, Ministry of Natural Resources, Xiamen 361005, China; Department of Microbiology, Biomedicine Discovery Institute, Monash University, Clayton, VIC 3800, Australia; Institute of Marine Biology and Pharmacology, Ocean College, Zhejiang University, Zhoushan 316021, China; Ocean Research Center of Zhoushan, Zhejiang University, Zhoushan 316021, China; State Key Laboratory for Marine Environmental Science, Institute of Marine Microbes and Ecospheres, College of Ocean and Earth Sciences, Xiamen University, Xiang’an Campus, Xiang’an South Road, Xiamen 361102, China

## Abstract

Deep-sea cold seeps host high microbial biomass and biodiversity that thrive on hydrocarbon and inorganic compound seepage, exhibiting diverse ecological functions and unique genetic resources. However, potential health risks from pathogenic or antibiotic-resistant microorganisms in these environments remain largely overlooked, especially during resource exploitation and laboratory research. Here, we analyzed 165 metagenomes and 33 metatranscriptomes from 16 global cold seep sites to investigate the diversity and distribution of virulence factors (VFs), antibiotic resistance genes (ARGs), and mobile genetic elements (MGEs). A total of 2,353 VFs are retrieved in 689 MAGs, primarily associated with non-pathogenic functions like adherence. Additionally, cold seeps harbor nearly 100,000 ARGs, as important reservoirs, with high-risk ARGs presenting at low abundance. Compared to other environments, microorganisms in cold seeps exhibit substantial differences in VF and ARG counts, with potential horizontal gene transfer facilitating their spread. These virulome and resistome profiles provide valuable insights into the evolutionary and ecological implications of pathogenicity and antibiotic resistance in extreme deep-sea ecosystems. Collectively, these results indicate that cold seep sediments pose minimal public health risks, shedding lights on environmental safety in deep-sea resource exploitation and research.

**Importance:** Understanding pathogenicity and antibiotic resistance in environmental reservoirs like deep-sea cold seeps is critical for assessing public health risks, particularly with increasing human activities such as deep-sea mining. This study offers the first comprehensive analysis of virulome, resistome, and mobilome profiles in cold seep microbial communities. While cold seeps act as reservoirs for diverse ARGs, high-risk ARGs are rare, and most VFs contribute to non-pathogenic ecological functions. These results highlight the minimal threat to public health posed by cold seeps, providing a reference for monitoring the spread of pathogenicity and resistance in extreme ecosystems and informing environmental safety assessments during deep-sea resource exploitation.

## Introduction

The deep-sea environment, defined as seawaters and sediments below 200 meters, constitutes the largest habitat on Earth, covering approximately 65% of the planet’s surface (1). Ensuring public safety in these vast and largely unexplored regions is of paramount importance, especially with the proposed development of deep-sea mining activities. The concept of “One Health”, first introduced in scientific literature about a decade ago (2), is now defined as an approach that “recognizes the health of humans, domestic and wild animals, plants, and the wider environment (including ecosystems) are closely linked and interdependent” (3). Therefore, it has become increasingly important to evaluate virulent and resistant risks not only within traditional medical or clinical settings but also to investigate potential environmental reservoirs where resistant and virulent bacteria may reside, evolve, and potentially be transmitted back to humans or animals.

Cold seeps, predominantly located along continental margins, are rich in gas hydrates, a type of deep-sea minerals(4). Gas hydrates are crystalline solids formed from water and gas, and they possess high commercial exploitation values (5). These seepage zones are formed by the upward migration of cold, hydrocarbon-rich fluids (mainly methane) through microfractures in the seafloor (6, 7). These locations often feature carbonate deposits and host abundant, diverse microbial population, which are supported by chemosynthetic bacteria, such as anaerobic methane-oxidizing archaea (ANME) that live in syntrophic associations with sulfate-reducing bacteria (SRB) (8, 9). Cold seep ecosystems, known as deep-sea oasis (9), are extraordinary deep-sea habitats with high biodiversity, offering a perspective for addressing deep-sea public safety concerns. However, for a long time, scientific research has focused on community diversity, ecological functions (such as biogeochemical cycling), evolutionary processes, and microbial genetic resources (4, 10–15). Little attention has been paid to the potential presence of pathogenic or antibiotic-resistant microorganisms in cold seeps.

The spread of infectious diseases caused by pathogenic bacteria is one of the leading causes of mortality in humans and animals (16, 17), and this issue has become increasingly unpredictable due to the rise of antibiotic-resistant bacteria (18). The pathogenesis associated with bacterial infections can be complex, but in many cases, it is primarily driven by virulence factors (VFs), produced by bacteria that enable them to persist, grow, and cause damage within the tissues of human or animal hosts (16, 19, 20). *Escherichia coli* and *Vibrio cholerae* are generally harmless to humans, but the acquisition of one or more genes can transform them into deadly pathogens (21, 22). It is surprising to discover that in the Mariana Trench, several well-known toxins such as botulinum neurotoxins, tetanus neurotoxin, and large clostridial toxins, have been detected in sediment samples from the Challenger Deep at a depth of 10,898 meters (23).

Antimicrobial resistance (AMR) has been recognized as a severe global public health threat (24–26). In fact, long before the large-scale production of antibiotics for the prevention and treatment of infectious diseases, many bacterial species had already evolved resistance to antibiotics, as seen in isolated caves (27, 28), permafrost cores (29) and human paleofeces (30). Beyond ancient environments, the environmental microbiome has been demonstrated to be a potential reservoir of antibiotic resistance genes (ARGs), which have emerged as a new class of contaminants across various ecosystems, including soil (31), aquatic environments (32–34), seawater (35–37), air (38), the human gut (39), wastewater treatment plants (40), and urban environments (41).

Amid selective pressure, interactions (synergistic selection) occur between ARGs and VFs (42, 43), which provides pathogens with greater advantages (44). VFs are essential for bacteria to overcome host defense systems, while the acquisition of antibiotic resistance enables bacteria to withstand antimicrobial therapies and adapt to and colonize extreme environments (45), posing a great threat to public health (46, 47). Moreover, mobile genetic elements (MGEs) such as integrons, transposons, and plasmids play a crucial role in the horizontal transfer of ARGs within microbial communities (47–49), leading to an increase in the number of antibiotic-resistant bacteria (ARB) (50, 51), which potentially enables the horizontal gene transfer of ARGs from environmental reservoirs to clinical pathogens (52).Therefore, investigating the pathogenic and antibiotic resistant potentials of microbial populations, is crucial for therapeutic strategies and biodefense analysis. In recent years, the types of the virulome and resistome across various ecosystems have been explored by various methods (20, 53–55), but these in cold seeps remain largely uninvestigated. In the One Health era (51), assessing the pathogenicity and antibiotic resistance of cold seep microorganisms provides valuable insights for the safety evaluation of future cold seep exploration, gas hydrates mining, sampling, and laboratory cultivation processes, guiding interventions to limit their spread.

In this study, we conducted a comprehensive analysis of the types and distribution of VFs, ARGs, and MGEs in microbial communities from 165 sediment samples collected across 16 globally-distributed cold seep sites. The counts of VFs, ARGs, and MGEs in Earth’s microbial genomes were also incorporated for comparative analysis (56). By analyzing these data, we identified 13 VF classes from 689 metagenome-assembled genomes (MAGs), with low abundance of toxin and invasion genes. We also detected numerous and diverse ARGs in cold seep sediments, which serve as a reservoir of ARGs. Moreover, the risk of these vast ARGs were assessed. To further investigate the potential for horizontal gene transfer of VFs and ARGs, we examined the diversity and distribution of MGEs. These findings provide the first systematic understanding of the pathogenicity and antibiotic resistance potential of deep-sea cold seep microbial communities, shedding lights on the assessment of public health and safety in deep sea.

## Results and discussion

### Fourteen classes of virulence factors were detected in cold seeps, which mainly facilitate adherence, serving non-pathogenic ecological roles

Fourteen classes comprising 14,023 VFs were identified in the non-redundant gene catalog from global cold seep sediments **(Fig. 1A)**, including all classes listed in VFDB. These VFs were primarily associated with ecological functions such as adherence, immune modulation, effector delivery systems, nutritional/metabolic factors and motility (**Fig. 1A**). Among them, adherence were the most abundant (28.59%), closely followed by those involved in immune modulation (28.48%). These results suggest that microorganisms in cold seeps rely heavily on adherence and motility for their ecological functions, likely aiding in surface attachment and navigation within the sediments (57, 58). Additionally, immune modulation genes potentially facilitate the establishment of symbiotic relationships with the seep fauna or to evade their immune responses and promote infection (e.g., infecting the nuclei of deep-sea mussels) (59, 60).

**Fig. 1.**
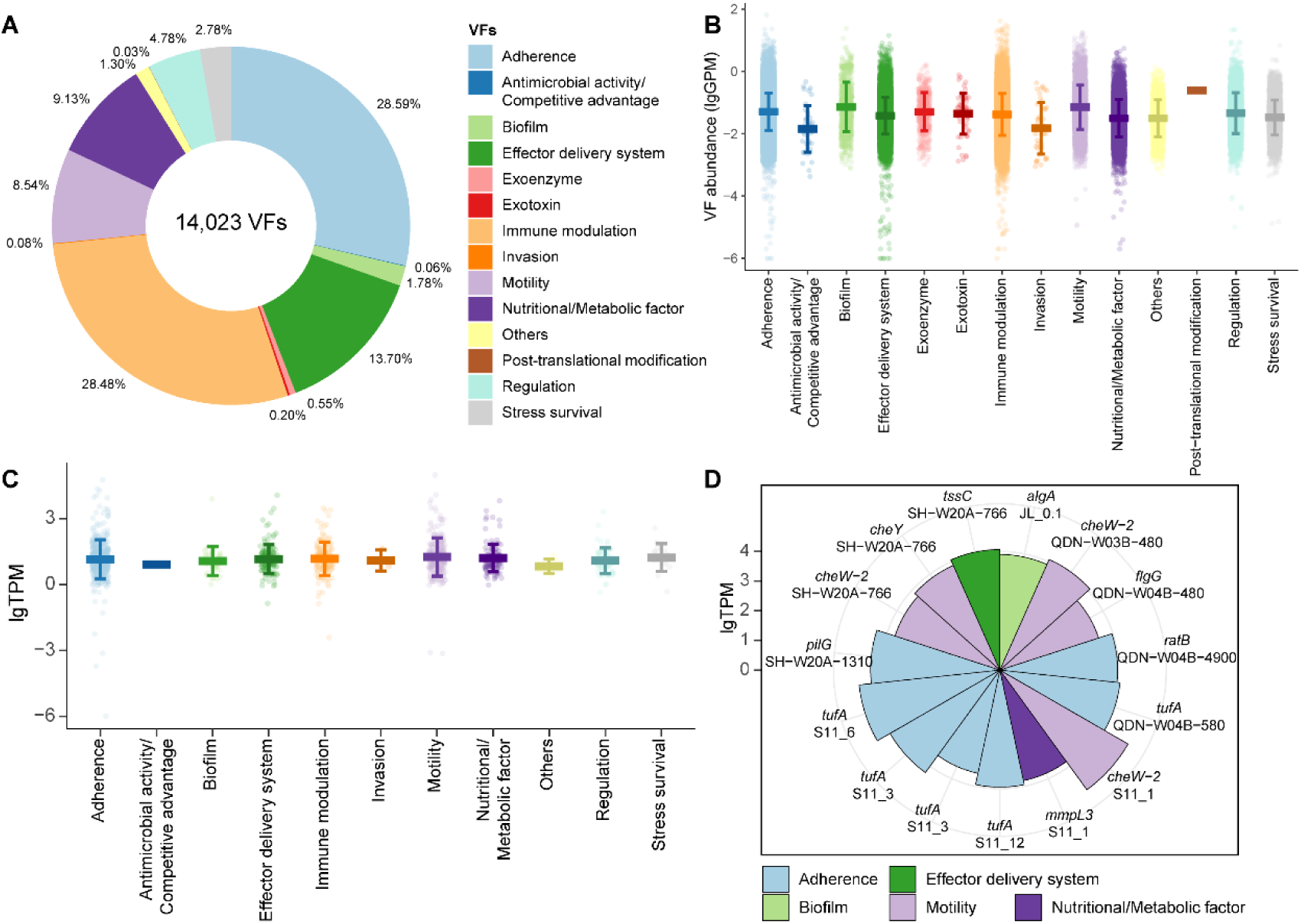
VFs detected in the non-redundant gene catalog from cold seep sediments. **(A)** Relative proportions of different VF categories in the non-redundant gene catalog, with each category represented by a distinct color as shown in the legend. **(B)** Abundances of VF categories at different cold seep sites. Each point represents the abundance of a VF gene at a specific site, with vertical bars denoting the minimum and maximum values. Gene abundances are expressed in Genes Per Million (GPM) and are plotted on a log10(x+1) scale. Details are shown in **Table S2**. **(C)** Transcript abundance of different categories of VFs from 33 cold seep sediment samples. Each point represents the transcript abundance of a VF gene at a cold seep site. Vertical bars indicate the minimum and maximum VF transcript abundances. Transcript abundances are represented in units of transcripts per million (TPM), with values shown on the graph as log10(x+1). Details are shown in **Table S3**. **(D)** Wind rose diagram showing the top 15 expression levels for different VFs in cold seeps. Transcript abundances are represented in units of transcripts per million (TPM), with values shown on the graph as log10(x+1). Each italicized text indicates a VF gene, with site abbreviation shown below each label.

Adherence VFs were widely distributed **(Fig. 1B and Table S2)** and highly expressed **(Fig. 1C and Table S3)** in cold seep sediments. These genes spanned multiple bacterial phyla **(Fig. S2)**, such as *tuf* and *tufA* (**Fig. 1D**, surface-expressed elongation factor Tu, EF-Tu), which mediate attachment by interacting with host cell nucleolin (61). Other highly expressed adherence genes included *pilG* (up to 22,508 TPM), which promotes twitching motility that allows bacteria to move along the cell surface (62), and *ratB* (up to 9.205 TPM), a putative outer membrane protein involved in intestinal colonization and persistence (63). These adherence genes probably assist microbes in stabilizing their ecological niches in cold seeps, thereby enhancing survival and nutrient acquisition.

In addition to adherence VFs, other non-pathogenic VFs involved in basic microbial physiological functions were identified, including effector delivery system (13.70%), nutritional/metabolic factors (9.13%), and motility (8.54%). Specifically, the effector delivery system gene *tssC* of type VI secretion system was detected at three cold seep sites, with expression levels reaching up to 11,529 TPM. Meanwhile, the nutritional/metabolic factor *mmpl3*, an MMPL family transporter, was expressed in two sites, with levels up to 6,424 TPM **(Fig. 1C)**. Among motility VFs, highly expressed genes included those related to chemotaxis proteins, such as CheW-2 (up to 95,590 TPM) and CheY (up to 7,216 TPM), as well as flagella proteins, such as FlgG (up to 3,126 TPM) **(Fig. 1D)**. This suggests that a suite of motility genes is required for navigating toward optimal living conditions, locating hosts or symbiotic partners (64, 65), and responding to the instability and limited resources in cold seep environments.

The remaining VFs associated with antimicrobial activity/competitive advantage, exotoxins, and invasion were detected at low abundances, with the mean abundance of 0.056, 0.104, and 0.067 GPM, respectively **(Fig. 1B and Table S2)**. Among these, 11 exotoxin genes (0.2%) were identified, including potentially harmful toxins such as the macrolide toxin mycolactone (66), and the pore-forming cholesterol-dependent cytolysin suilysin (67). However, none of these exotoxin genes were expressed at any cold seep sites **(Table S3)**. Aside from exotoxins, exoenzyme and post-translational modification genes also showed no expression, whereas the other 11 VF categories exhibited transcriptional activity **(Fig. 1D)**. For the invasion-related genes, only *ibeC* displayed low transcriptional level, reaching 44.76 TPM in the Jiaolong cold seep **(Fig. 1D and Table S3)**, a gene known to facilitate the invasion of brain microvascular endothelial cells (68). The low abundance and lack of expression of pathogenic genes indicate that the potential threat to human health from global cold seeps is minimal.

### Cold seep virulome are enriched in *Actinomycetota* and *Pseudomonadota* phyla

Among the 3,164 species-level MAGs recovered from cold seep sediments, 689 MAGs were identified with 13 classes of VFs (n = 2,353), spanning 68 bacterial phyla (n = 2,343) and five archaeal phyla (n = 10) **(Figs. 2A-C and Table S4)**. In bacteria, the majority of VFs (42.85%) were related to adherence, followed by motility (18.44%) and immune modulation (11.44%) (**Fig. 2B**). In archaea, 10 VFs were associated with immune modulation, with these factors detected across several archaeal phyla, including *Thermoplasmatota*, *Nanoarchaeota*, and *Asgardarchaeota*. Notably, *Thermoplasmatota* accounted for 40% of all identified archaeal VFs (**Fig. 2C**).

**Fig. 2.**
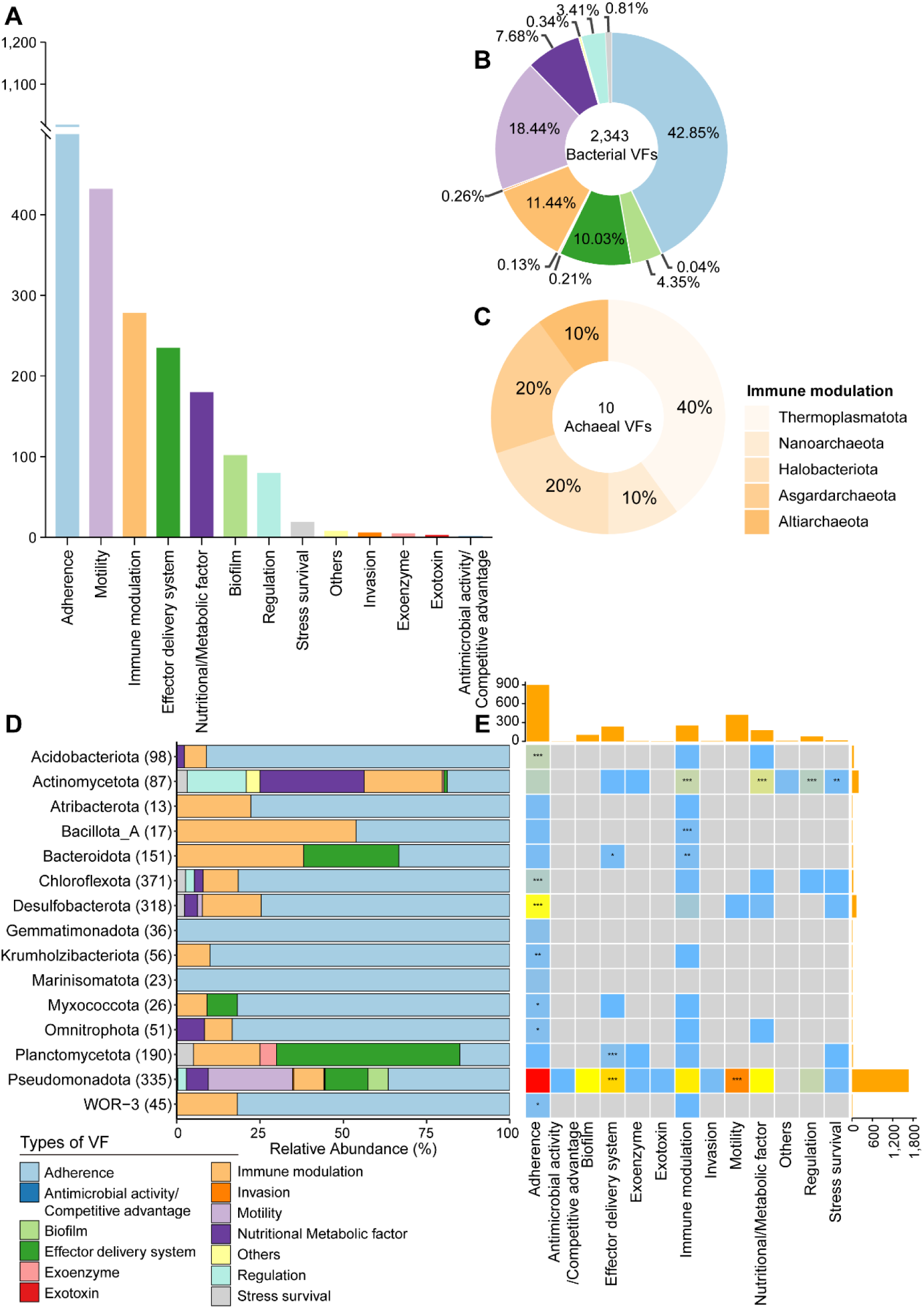
Taxonomic distribution of VFs. **(A)** Bar chart showing the counts of different VF categories identified in archaeal and bacterial MAGs derived from 165 cold seep sediment samples. Additional details are provided in **Table S4**. **(B)** Relative proportions of different VF categories across bacterial phyla. **(C)** Relative proportions of archaeal phyla within immune modulation VFs. **(D)** Relative proportions of VF categories across the top 15 taxonomic phyla. The number of MAGs in each phylogenetic cluster is indicated in brackets. Different VF categories are colored as indicated in the legend. **(E)** Association of VF categories with different phylum. Asterisks indicate significant enrichment according to Fisher’s exact test (*/**/***, Odds ratio>1 and P-value < 0.05/0.01/0.001 after Benjamini-Hochberg correction). The heatmap colors exhibit the number of each VF category in each phylum. The bar chart at the top represents the total count of each VF category across all the 15 phyla, while the bar chart on the right shows the total count of all VF categories for each phylum.

The enrichment of different categories of VFs across the top 15 microbial phyla was analyzed using Fisher’s exact test, with significance defined as an odds ratio > 1 and p < 0.05 **(Fig. 2D and Fig. 2E)**. Although both *Actinomycetota* and *Pseudomonadota* had abundant and diverse VFs, the dominant types differed between these two phyla. In *Actinomycetota*, nutritional/metabolic factor genes were significantly enriched, such as *panD* (aspartate-1-decarboxylase), involved in pantothenate biosynthesis (69) and *ctpC*, a P(1)-type Mn^2+^ transporting ATPase. These genes are essential for microbial growth and environmental adaptation. *Actinomycetota* also showed significant enrichment in immune modulation genes, primarily *acpXL*, an acyl carrier protein, and *gmd* (GDP-mannose 4,6-dehydratase), assisting in evading the host immune system. In contrast, *Pseudomonadota* exhibited significant enrichment in VFs related to motility and effector delivery systems. Motility genes primarily included polar and peritrichous flagella, while effector delivery genes were largely associated with types II, III, and VI secretion systems. These systems promote the translocation of proteins or toxins from the bacterial cell to the extracellular environment or directly into host cells (70).

Notably, 1,196 VFs were recognized in 17 MAGs, accounting for more than 50% of the total, primarily belonging to *Actinomycetota* and *Pseudomonadota* phyla **(Fig. S3)**. Among these, the VFs in *Escherichia coli* (5.8%) and *Salmonella enterica* (10.5%) within the *Pseudomonadota* phylum were probably attributed to contamination. *Escherichia coli* correlate with experimental contaminant (71), while *Salmonella* is often introduced into marine environments through rivers or stormwater (72, 73). In cold seeps, *Salmonella enterica* may have contaminated sediments via filter-feeding mussels (73).

Other VFs-enriched microbial groups in cold seeps, such as *Vibrio diabolicus* (n = 196), *Pseudomonas_E sp002874965* (n = 139), and *Stutzerimonas frequens* (n = 85), contained one or none of the key exotoxin/invasion genes, as previously mentioned **(Fig. S4)**, in contrast to those found in the Mariana Trench that encode toxins known to affect human health (23). Similar to other habitats (e.g., global natural ecosystems and host-associated environments), most VF-enriched microbes were affiliated with the phyla *Pseudomonadota* (formerly *Gammaproteobacteria*), particularly from *Enterobacteriaceae* and *Pseudomonadaceae* families (56). Some VFs-enriched microbial groups in cold seeps were considered to be source-related. For example, *Vibrio diabolicus*, a deep-sea facultative anaerobic heterotroph isolated from the polychaete *Alvinella pompejana* in the East Pacific hydrothermal vent field (74), has a close relationship with its annelid host. This potential host-associated origin explains the abundance of adherence, motility, effector delivery systems, and immune modulation genes in its genome.

### Cold seep archaeal and bacterial phyla harbor nearly one hundred thousand antibiotic resistance genes

Combining annotation results from a range of databases, 92,532 ARGs were identified across 3,163 MAGs, representing 16 archaeal and 99 bacterial phyla **(Table S5)**. Only one MAG (SH-W20A-3880_sbin_30), belonging to the archaeal phylum *Nanoarchaeota*, was inferred without any ARGs. In total, 27 types of ARGs were retrieved in cold seep MAGs, with four types of multidrug, aminoglycoside, MLS (macrolide-lincosamide-streptogramin), and beta-lactam resistance genes exceeding 10,000 counts **(Fig. 3A)**. In archaea, 9,576 ARGs belonging to 25 types were detected, with multidrug resistance (MDR, 34.49%) and aminoglycoside resistance (28.41%) being the most prevalent, followed by MLS (17.20%) and beta-lactam (8.71%) resistance **(Fig. 3B)**. In contrast, bacterial ARGs were more diverse, comprising 82,956 genes across 27 types, where MDR and aminoglycoside resistance genes made up 33.09% and 23.36%, respectively **(Fig. 3C)**. These patterns suggest distinct resistance strategies between archaea and bacteria in cold seeps.

**Fig. 3.**
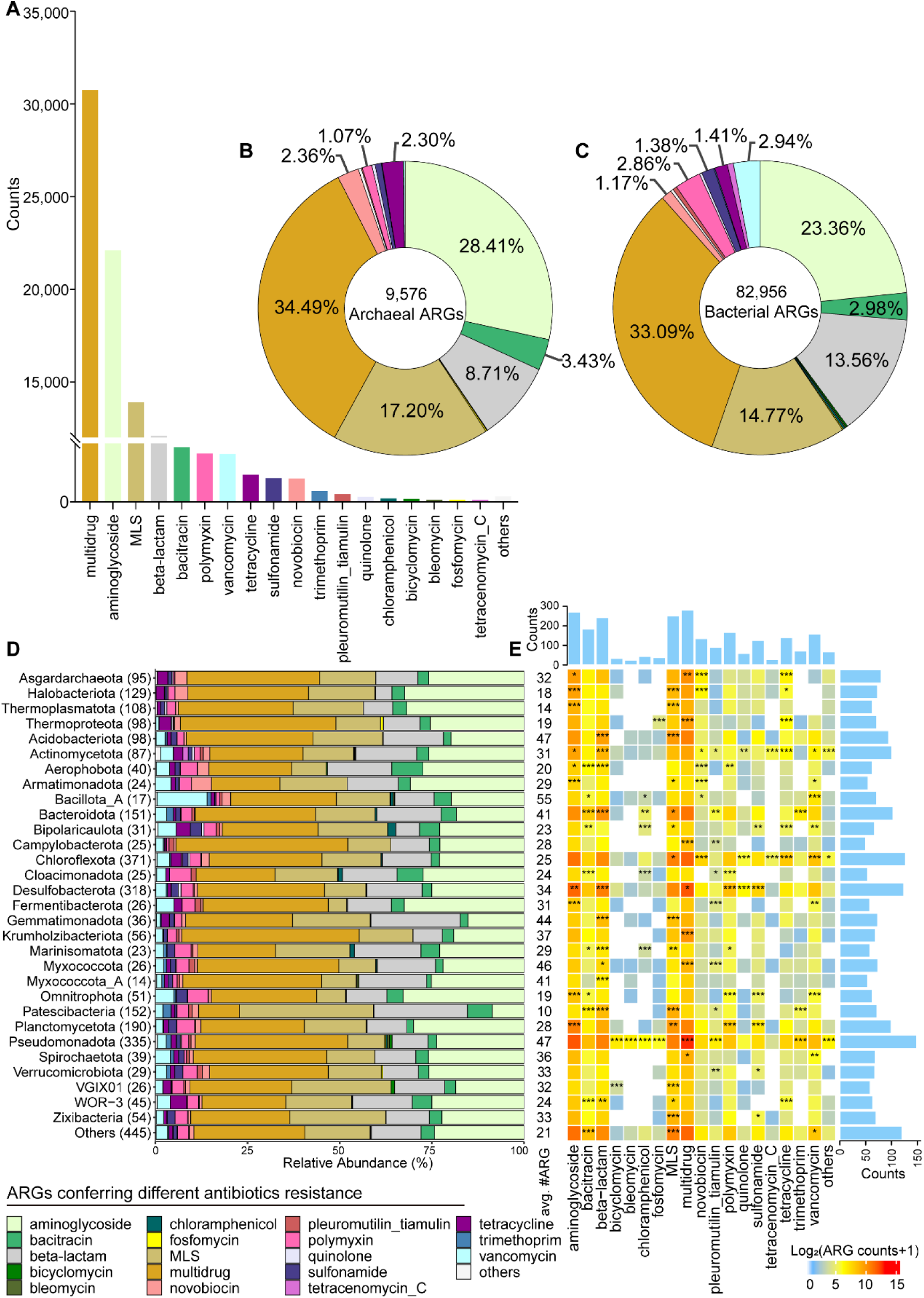
Different ARGs across phyla and sites in cold seeps. **(A)** Bar chart showing the counts of ARGs conferring different antibiotics resistance identified in archaeal and bacterial MAGs derived from 165 cold seep sediment samples. Details are shown in **Table S5**. **(B-C)** Relative proportions of different ARG types across **(B)** archaeal and **(C)** bacterial phyla. **(D)** Relative abundance of ARG types across different microbial phyla (ARGs > 500). The number of MAGs in each phylogenetic cluster is indicated in brackets. Different ARG types are colored as indicated in the legend. **(E)** Association of ARG types with different phyla. Asterisks indicate significant enrichment according to one sided Fisher’s exact test (*/**/***, Odds ratio>1 and P-value < 0.05/0.01/0.001 after Benjamini-Hochberg correction). The heatmap colors exhibit the number of each ARG type in each phylum, log-transformed (Log2(x + 1)) for plotting. The bar chart at the top represents the total count of each ARG type across all phyla, while the bar chart on the right shows the total count of all ARG types for each phylum.

The enrichment of various ARGs in different microbial phyla (ARGs > 500) was analyzed using Fisher’s exact test, with significance defined as an odds ratio > 1 and p < 0.05 **(Figs. 3D-E)**. Among archaea, the phyla *Asgardarchaeota*, *Halobacteriota*, *Thermoplasmatota*, and *Thermoproteota* exhibited a high diversity of ARG types, with the *Asgardarchaeota* phylum showing the highest average count of ARGs per genome (n = 32). In bacteria, ARGs were abundant and diverse in the phyla *Pseudomonadota*, *Chloroflexota*, and *Actinomycetota*, while *Bacillota_A* had the highest average count of ARGs per genome (n = 55). MLS resistance and MDR genes were significantly enriched in multiple bacterial and archaeal phyla.

Not all ARGs pose serious threats to public health, so a proposed omics-based framework was applied to identify those risk ARGs in cold seeps by analyzing enrichment of ARGs in anthropogenically-impacted environments, mobility, and host-pathogenicity (75). High-risk ARGs in cold seeps, including current (Rank I, n = 9.63%) and future (Rank II, n = 1.59%) threats to human health, showed low abundance across cold seep sites **(Fig. S5)**, accounting for 11.22% of the total abundance of Rank I-IV ARGs. Rank III (human-associated and non-mobile) ARGs accounted for 11.26% in cold seeps, while the remaining 77.51% were Rank IV ARGs. In contrast to global wastewater microbiomes, where Rank I and II ARGs account for 67.0% of the total abundance (76), the cold seep microbiome exhibits a minimal potential public health risk posed by ARGs.

### The abundance and transcript level of antibiotic resistance genes vary across different cold seep sites

The abundance of different ARG types varied across cold seep sites **(Fig. S6 and Table S6)**. A total of 812 ARG subtypes were annotated based on metagenomic reads using ARGs-OAP v3.2.4, from which the top 100 abundant ARG subtypes were selected for display **(Fig. 4A)**. Among 812 ARG subtypes, the abundance of *PNGM-1*, *macB*, *arnA*, *ugd*, *ranA*, *ranB*, *msbA*, *bacA*, and *novA* genes were consistently high across different types and depths of global cold seeps. In most cold seep sites, beta-lactam resistance genes, with a mean abundance of 0.012 GP16S (gene copies per 16S rRNA gene), and multidrug resistance (MDR) genes, with a mean abundance of 0.006 GP16S, were the most abundant. Notably, the cold seep at the ENP site (Pacific Ocean: Eastern North Pacific, ODP site 1244) exhibited the highest overall abundance (sum of 4.656 GP16S), with aminoglycoside resistance genes being the primary contributors, reaching up to 1.011 GP16S. These genes are mainly associated with resistance to aminoglycoside antibiotics, such as streptomycin and spectinomycin, predominantly natural products produced by *Actinomycetota* (77). ENP cold seep is subject to high levels of aminoglycoside antibiotic selective pressure, potentially due to the presence of multiple Actinomycetotal groups producing these antibiotics. For example, *Hydromicrobium sp004376325* (QDN-W03B-2880_sbin_3) accounts for 4.33% in ENP_18.1 site, while the genus *Hydromicrobium* (ENP_cobin_7) accounted for 3.33% of ENP_18.1 and over 1% of ENP_2, ENP_35.65, and ENP_68.55 **(Table S7)**.

**Fig. 4.**
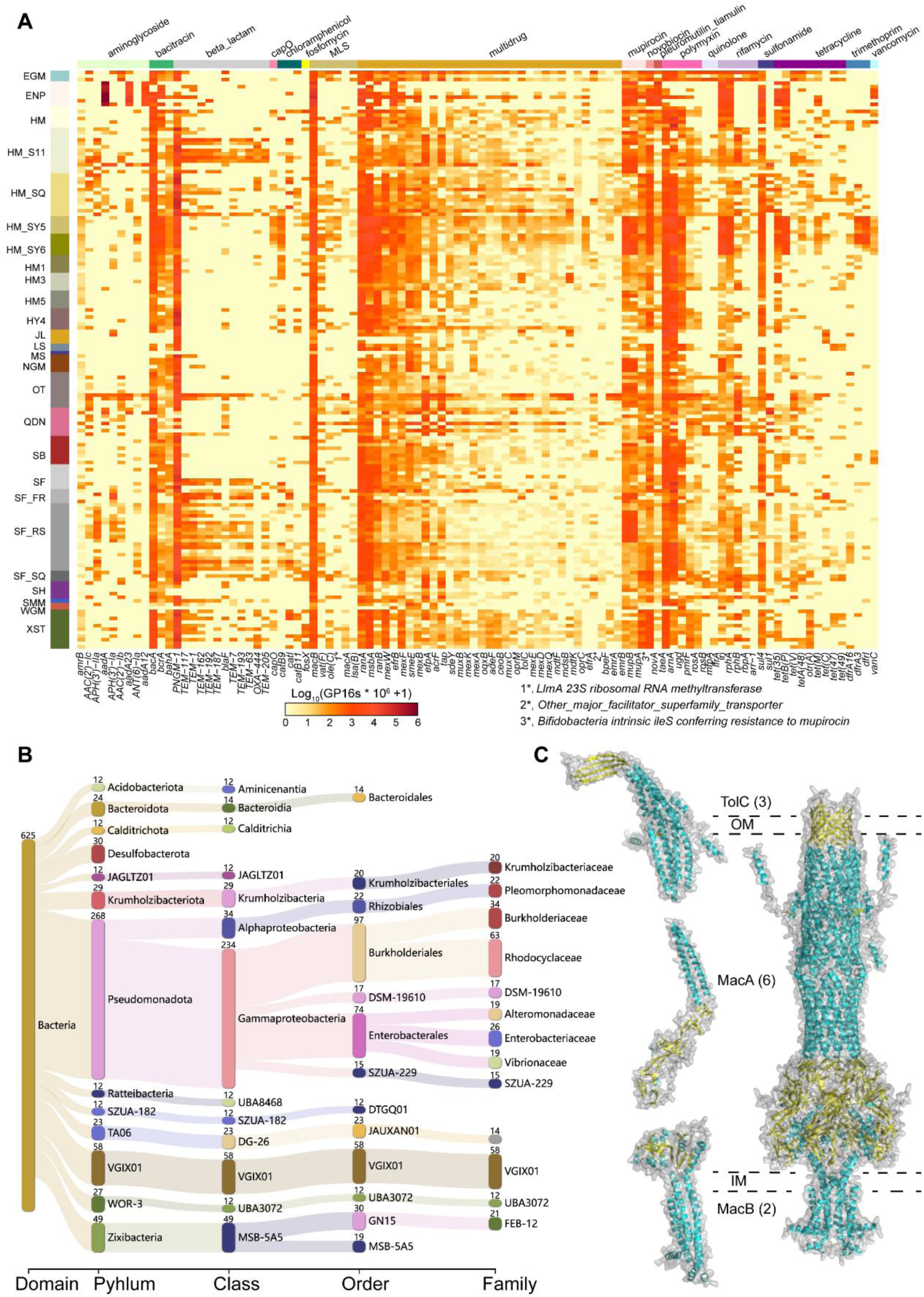
Deep-sea cold seep microbiomes harbor diverse efflux pumps conferring antibiotic resistance. **(A)** The heatmap displaying gene abundances of the top 100 abundant ARG subtypes across different cold seep sites. Abundance was log-transformed (i.e., Log10(GP16S × 10^6^ + 1)) for the plot. Each row indicates a sample (arranged by sample type). Each row represents a cold seep site, with sample abbreviations indicated on the left. Each column represents an ARG subtype, affiliated with an ARG type indicated by different colors at the top. **(B)** Sankey diagram illustrating the taxonomic distribution of MAGs containing the *macAB-tolC* pump. **(C)** Predicted structures of MacA, MacB, and TolC proteins, as well as their assembly into the complete efflux pump complex, predicted by AlphaFold3. The left side shows the individual protein structures, while the right side illustrates the predicted interactions, forming the trimeric TolC, hexameric MacA, and dimeric MacB complexes. The full efflux pump comprises 11 proteins, with outer membrane (OM) and inner membrane (IM) boundaries indicated by dashed lines.

Compared to other environments, the average abundance of ARGs in cold seeps was significantly lower than that in rivers upstream and downstream of sewage treatment plants (p < 0.001; **Fig. S7**), consistent with findings that ARG abundance increases with human activity (75, 76, 78, 79). Surprisingly, the average ARG abundance in cold seeps was also lower than in the abyssal Mariana Trench **(Fig. S7)**, despite the trench being far away from human activities. Due to the “V”-shaped structure of hadal trench acting as a natural collector of organic pollutants, the accumulation of persistent organic pollutants (e.g., PCBs, PBDEs, and PAHs) in trench sediments and fauna (80) likely promotes the selection of resistance genes in trench bacteria (81–83).

Metatranscriptomic data indicated that transcriptionally active ARGs conferred resistance to 27 classes of antibiotics **(Table S8)**. Various ARGs were expressed across different cold seep sites **(Fig. S8 and Table S8)**, with MDR accounting for 34.28% of all expressed ARGs, followed by aminoglycoside resistance (25.19%), MLS resistance (13.15%), and beta-lactam resistance (13.09%). Naturally occurring antimicrobial secondary metabolites (e.g., antibiotics) are widely recognized as mediators of competition between microbial species in both soil and marine environments (14, 84, 85). Consequently, these expressed ARGs in cold seep sediments play a crucial role in enabling microbes to evade the inhibitory or lethal effects of antibiotics.

### Diverse efflux pumps conferring antibiotic resistance are widely distributed in cold seep microbes, with ABC systems being predominant

Efflux pumps play a crucial role in microbial resistance to multiple antibiotics by expelling toxic compounds, including antibiotics, out of the cell. In cold seeps, MDR genes exhibited the great diversity and high abundance **(Fig. 4A)**. MDR is crucial for bacterial survival, especially through MDR pumps, which provide resistance against multiple classes of antibiotics and help bacteria endure competition with epibiotic bacteria and host-produced antimicrobials (86, 87). The most prevalent MDR pump system was the ATP-binding cassette (ABC), with *ranA* (up to 0.0197 GP16S) and *ranB* (up to 0.0061 GP16S) forming an ABC-type efflux system, followed by *msbA* (up to 0.0083 GP16S).

Besides ABC systems, cold seep microbes also harbor four other types of efflux systems: major facilitator superfamily (MFS), e.g., *qacEdelta1* (up to 0.0153 GP16S) and *efpA* (up to 0.0066 GP16S); multidrug and toxic compound extrusion (MATE), e.g., *mdtK* (up to 0.0060 GP16S) and *pmpM* (up to 0.0013 GP16S); small multidrug resistance (SMR) systems, e.g., *qacH* (up to 0.0007 GP16S) and *ykkD* (up to 0.0002 GP16S); and resistance-nodulation-cell division (RND) systems, e.g., *mexB* (up to 0.0095 GP16S), *mexA* (up to 0.0006 GP16S), and *oprM* (up to 0.0006 GP16S) **(Table S6)**. MDR pumps are not only important components for antibiotic resistance but also play a role in bacterial pathogenicity (86, 87). In addition to providing protection against host-produced antimicrobial compounds, they belong to regulatory networks that encompass VFs (86, 88, 89). MDR pumps in cold seeps are instrumental in mediating interactions between commensal and pathogenic microorganisms and their hosts.

Of ABC-type multidrug efflux systems in cold seep microbiome, *msbA*, besides *ranAB*, exhibited high gene abundance across global cold seep sites **(Fig. 4A)**. A total of 306 *msbA* genes were recognized, with archaea belonging to the phyla *Asgardarchaeota* (class *Heimdallarchaeia*, n = 11) and *Thermoproteota* (n = 2). Among bacteria, 49 phyla were annotated with *msbA*, with the high prevalence in *Pseudomonadota*, *Desulfobacterota*, *Bacteroidota*, and *Acidobacteriota* **(Fig. S9A)**. The MsbA proteins in cold seep microorganisms demonstrated close phylogenetic relationships to reference sequences and exhibited evolutionary conservation when compared to reference structures **(Fig. 5A, Fig. S9B and Table S9)**. MsbA functions as a homodimer, relying on dimeric interactions for its activity. The phylum *Desulfobacterota* serve as key players in cold seeps, and some species have sulfate-reducing potential. FR_cobin_47 belonged to the family *Desulfatiglandaceae* carries the dsrA gene (a key gene for sulfate reduction) (90, 91), and is present in more than 70% of cold seeps **(Table S7)**. Its MsbA dimer structure was predicted and aligned with the PDB reference structure (3B5Z) **(Fig. 5B)**, with the complexqTMscore (TM-score of complex alignment normalized by the query length) of 0.86982 and complextTMscore (TM-score of complex alignment normalized by the target length) of 0.93064. TM scores above 0.8 indicate a high degree of structural similarity, suggesting that the predicted dimeric structure of MsbA closely matches the experimentally crystal structure. This structural conservation reflects the evolutionary stability of MsbA’s functional domains, which include two characteristic coupling helices, two substrate-specific α-helical transmembrane domains (TMDs), two highly conserved nucleotide-binding domains (NBDs), and well-formed binding sites. This observation aligns with previous studies, which indicate that genes encoding multidrug efflux pumps are evolutionarily ancient and highly conserved elements (86). MDR pumps have other physiologically relevant roles such as detoxification of intracellular metabolites, intercellular signaling, stress response and cell homeostasis in the natural ecosystems (87, 92). Over billions of years, most ARGs likely evolved from genes with other original functions (93, 94). In antibiotic-producing bacteria, MDR pumps primarily detoxify intracellular antibiotics rather than provide resistance to external antibiotics (86). Thus, these efflux pumps potentially serve roles beyond antibiotic resistance, mainly regulating bacterial behavior in deep-sea cold seep sediments.

**Fig. 5.**
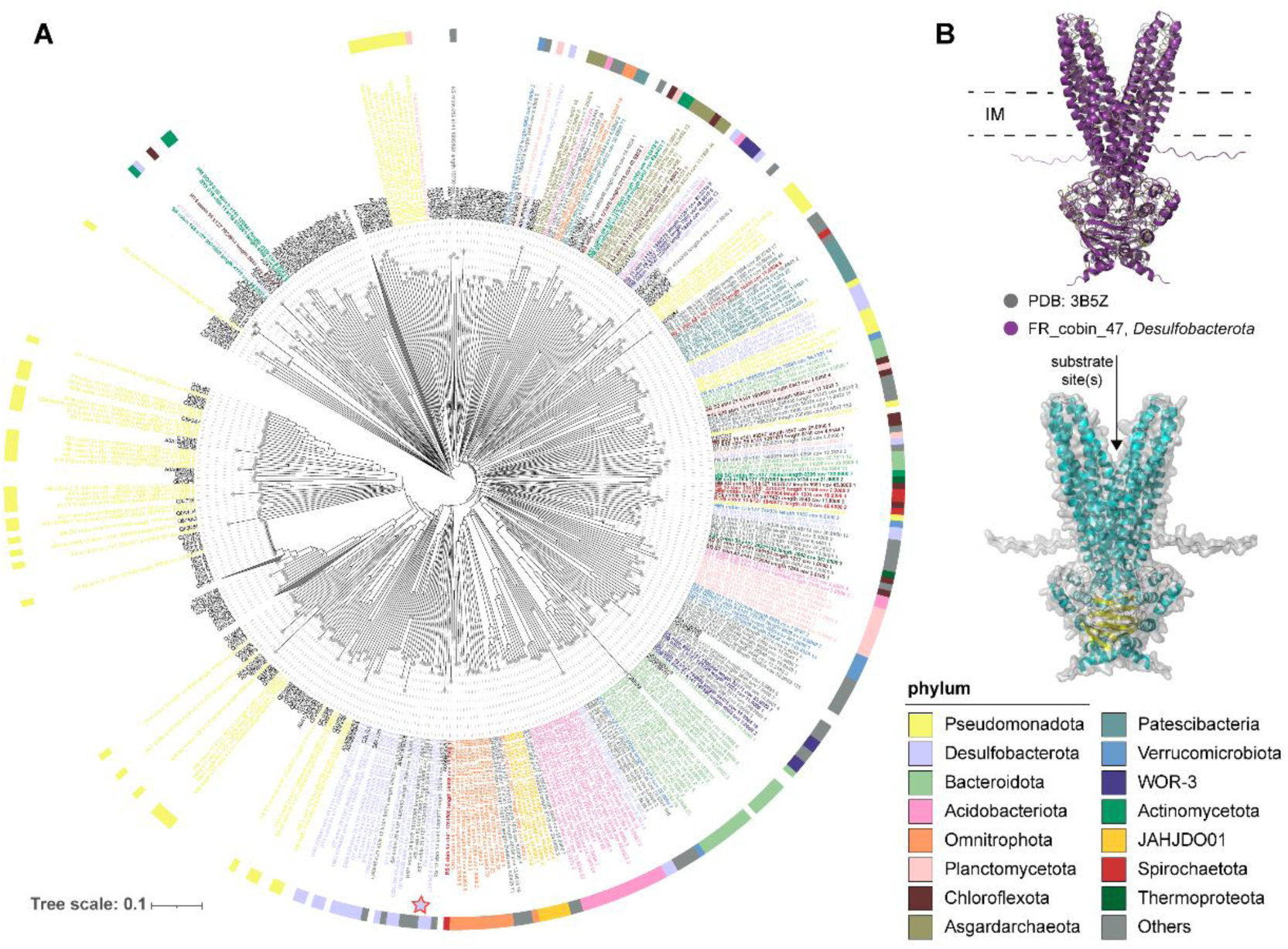
Tree and alignment of MsbA recovered from 3,164 cold seep MAGs. **(A)** Structural tree of MsbA predicted by ESMFold (n = 260) and the reference structures (n = 202) downloaded from the AlphaFold Protein Structure Database (AlphaFoldDB) and the Protein Data Bank (PDB). Reference structures are colored in black **(B)** Predicted structure of MsbA homodimer. The top section shows the alignment of predicted structure from FR_cobin_47 and the crystal structure of MsbA from *Salmonella typhimurium* (PDB ID: 3B5Z). The bottom section displays the predicted MsbA homodimer structure from FR_cobin_47, with inner membrane (IM) boundaries indicated by dashed lines.

MLS resistance genes were various (88 subtypes) and abundant (up to 0.0538 GP16S) as well **(Fig. 4A)**, The MacAB-TolC efflux system operates through the coordinated function of three components, 625 MAGs containing all three genes were selected for subsequent analysis, spanning 22 bacterial phyla **(Fig. 4B)**. Among these, *Pseudomonadota* was the most represented phylum (n = 268), followed by VGIX01 (n = 58), *Zixibacteria* (n = 49), and *Desulfobacterota* (n = 30). In these microbial groups, MacB forms a tripartite complex with the outer membrane protein TolC and the periplasmic partner protein MacA. The transport processes are coupled to ATP hydrolysis by MacB. The *macA*, *macB*, and *tolC* genes, along with their proteins predicted, exhibit close phylogenetic relationships to reference sequences and structures **(Fig. S10, Fig. S11 and Tables S10-12)**. The MacAB-TolC efflux pump of SZUA-182 MAG (SB_cobin_127) was further analyzed, which is distributed in multiple cold seeps including Eastern Gulf of Mexico, Scotian Basin, East China sea (Okinawa), South China Sea (Qiongdongnan Basin, Site F cold seep, and Haima) **(Table S7)**. The predicted MacAB-TolC efflux pump in the SB_cobin_127 was aligned with the PDB reference structure (5NIL, **Fig. S11D**), with the complexqTMscore of 0.85279 and complextTMscore of 0.81650, indicating high structural similarity to the experimentally determined crystal structure. In this structure, the periplasmic protein MacA forms a hexameric bridge between the trimeric TolC in the outer membrane and the dimeric MacB in the inner membrane, creating a quaternary structure with a central substrate translocation channel **(Fig. 4C)**. These pumps not only mediate the efflux of macrolide antibiotics but also transport outer membrane lipopeptides, protoporphyrin, polypeptide VFs, and lipopolysaccharides (92).

### Virulence factors and antibiotic resistance genes potentially spread in cold seeps through horizontal gene transfer

Secreted and non-secreted VFs were predicted using PathoFact, with localization determined across chromosomes, plasmids, phages, etc. Although most VFs were chromosomally encoded, 15% were located on plasmids or phages **(Fig. 6A)**, suggesting the potential for HGT of VFs in cold seeps. VFs on plasmids or phages were predominantly associated with members of the *Pseudomonadota*, *Chloroflexota*, *Asgardarchaeota*, and *Planctomycetota* phyla **(Fig. 6B)**. For these mobile VFs, the majority were located on plasmids in most phyla, such as *Hydrogenedentota* (34.29% of all VFs) and *Poribacteria* (32.50% of all VFs) **(Fig. S12)**, indicating that plasmid-mediated HGT is a key mechanism for the spread of VFs among cold seep microorganisms. The widespread transfer of plasmid DNA requires physical contact between microbes or between microbes and plasmids (95), which may be facilitated by the upward migration of cold, hydrocarbon-rich fluids (mainly methane) through microfractures in the seafloor, creating opportunities for physical contact of microbes in cold seep sediments. In some phyla, mobile VFs were primarily located on phages, such as *Bacillota_I* (19.17% of all VFs) and *Asgardarchaeota* (16.98%), **(Fig. S12)**, which enables genetic exchange over greater distances between bacteria. In cold seeps, as a subsurface reservoir of viral diversity (96), phage-mediated HGT contributes to the spread of VFs, potentially impacting bacterial pathogenicity.

**Fig. 6.**
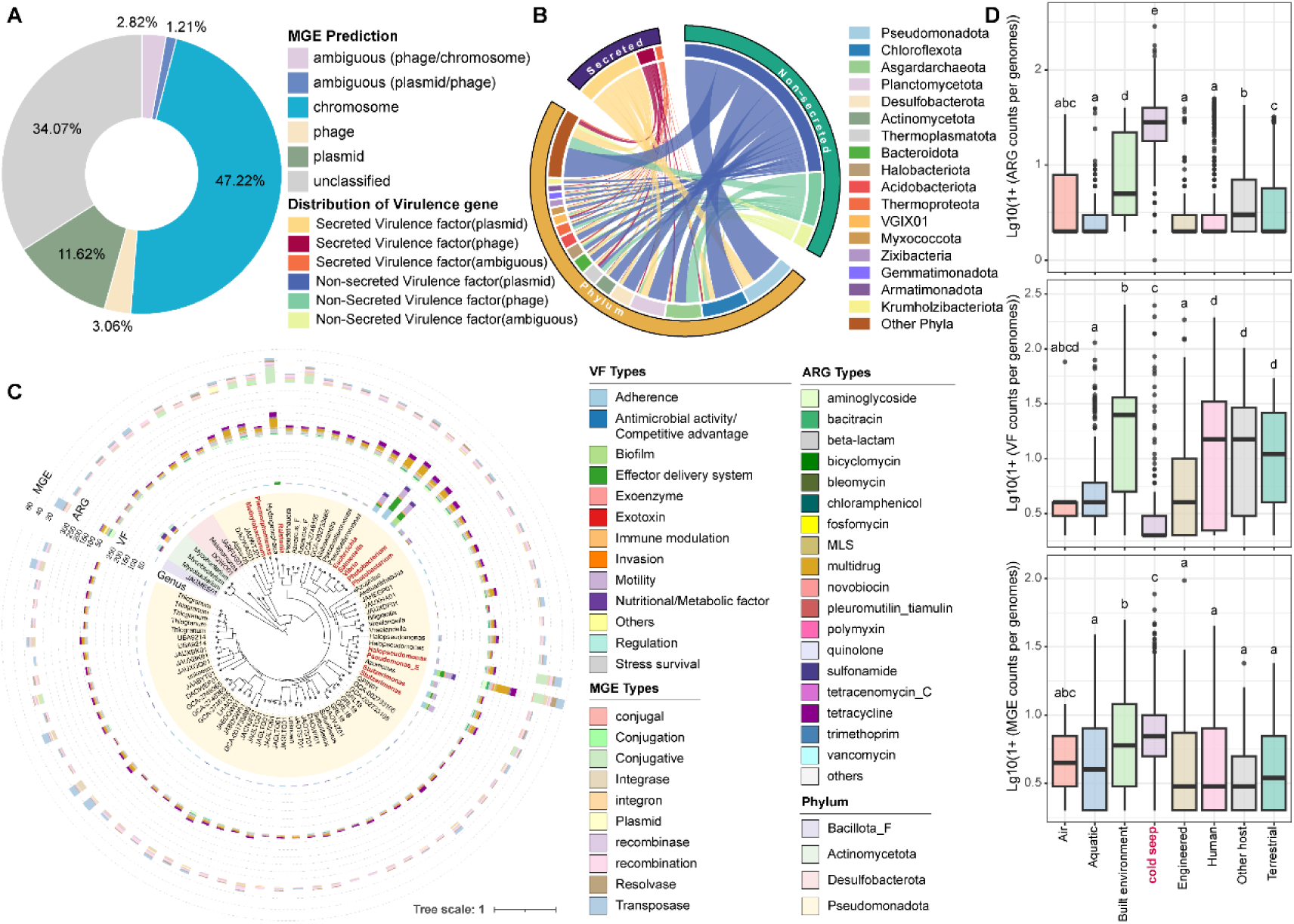
Comparison of ARGs, VFs, and MGEs identified in cold seep sediments. **(A)** Relative proportion of different VF positions detected in 3,164 cold seep MAGs. The positions of VFs are categorized into different types, including ambiguous (phage/chromosome), ambiguous (plasmid/phage), chromosome, phage, plasmid, and unclassified. **(B)** A chord diagram illustrating the association between different microbial phyla and the types of VFs carried on MGEs. **(C)** Phylogenetic tree of 82 VF-ARG carrying MAGs identified in cold seep sediments. The outer rings display the types and quantities of VFs, ARGs, and MGEs present in these genomes. **(D)** Boxplots comparing the counts of ARGs, VFs, and MGEs per genome across different ecosystems, including cold seeps and other habitats.

Out of 3,164 MAGs, 689 were identified as carrying both VFs and ARGs, accounting for approximately 21.8%. MGEs were detected in all these MAGs, except for OT_A2_sbin_55. Among them, 82 MAGs contained more than three VFs, ARGs, and MGEs, belonging to four bacterial phyla **(Fig. 6C)**. In these VF-ARG carrying MAGs, most VFs were related to adherence, effector delivery systems, and motility, while most ARGs conferred resistance to beta-lactam, multidrug, and aminoglycoside. MGEs were predominantly of the conjugative, recombination, and transposase types. Additionally, the co-localization of VFs and ARGs on contigs revealed their upstream and downstream proximity, with neighboring genes often containing MGEs **(Fig. S13)**, suggesting the co-existence of ARGs and VFs within pathogenicity islands (PAIs). PAIs, typically located in bacterial genomes, are transferred through horizontal gene transfer events such as plasmids, phages, or conjugative transposons, facilitating microbial evolution (97). The transfer of PAIs in cold seeps may stimulate the spread of virulence and resistance genes within microbial communities, potentially converting non-pathogenic bacteria into pathogens or conferring acquired antibiotic resistance, thereby reshaping microbial populations and community dynamics.

Comparing the counts of VFs, ARGs, and MGEs in cold seep microbial genomes with those in other habitats (56), we observed that the counts of VFs in cold seep sediments are significantly lower than in other global ecosystems, except for air **(Fig. 6D)**, indicating there are few pathogenic microorganisms in cold seeps. By contrast, the counts of ARGs and MGEs carried in cold seep microorganisms are significantly higher compared to other habitats **(Fig. 6D)**. This suggests that antibiotic resistance in deep-sea cold seeps is likely driven by HGT mediated by plasmids, transposons, or integrons, leading to the rapid spread and accumulation of ARGs within microbial communities. Antibiotics and common biocides stimulate the horizontal transfer of resistance (98, 99), and many naturally occurring stress can also accelerate HGT (100). Thus, the extreme environmental pressures in cold seeps, along with the selective pressure from microbes-produced antimicrobial compounds, possibly stimulate HGT and enhance the spread of ARGs, reflecting unique adaptive traits of microorganisms inhabiting the sediments of cold seeps.

In this study, we depict the landscape of virulome, antibiotic resistome, and mobilome in deep-sea cold seep sediments. The results generally indicated that VFs contribute to adherence and motility, playing a crucial role in non-pathogenic ecological functions. A few specific microbes from *Actinomycetota* and *Pseudomonadota* were VF-enriched taxonomic groups, accounting for over half of the identified VFs, such as *Vibrio diabolicus* and *Pseudomonas_E sp002874965*. Moreover, cold seeps serve as reservoirs for ARGs, with diverse efflux pumps being the predominant type, while MDR pumps are essential for mediating interactions between commensal and pathogenic microorganisms and their hosts. Although ARGs retrieved in cold seeps were diverse, high-risk ARGs were found to be of low abundance across global sites. Compared to other ecosystems, microbiome in cold seeps exhibited distinct signatures in VFs and ARGs, potentially transferred through horizontal gene transfer mechanisms like plasmids, phages, or conjugative transposons. However, understanding the evolutionary and ecological impacts of pathogenicity and antibiotic resistance in natural environments remains challenging due to the complex interplay of evolutionary, ecological, and environmental factors. Overall, these results demonstrate cold seep sediments pose minimal risks to public health. The resistome and virulome profiles provided here can serve as a reference for monitoring the evolution and spread of pathogenicity and antibiotic resistance in deep sea, while also offering foundational insights to guide future public health and safety assessments in these ecosystems, especially as human activities increase.

## Materials and methods

### Metagenomic and metatranscriptomic datasets

Metagenomic datasets were collected from 165 samples across 16 cold seep sites worldwide, and 33 metatranscriptomes were obtained from four of these sites. Non-redundant gene and genome catalogs were constructed following the methodology described in our previous study (101). Briefly, raw reads were trimmed to generate clean reads, which were then assembled into contigs. Protein-coding sequences were predicted and clustered to form a non-redundant gene catalog containing 147,289,169 representative clusters. Contigs longer than 1,000 bp were selected for binning, resulting in MAGs that were dereplicated at 95% average nucleotide identity (ANI). This process yielded 3,164 representative MAGs (each at least 50% complete and with less than 10% contamination) at the species level, which were taxonomically annotated using GTDB-Tk (v2.4.0) (102) against Genome Taxonomy Database GTDB (release 09-RS220) **(Table S1)**. The coverage of each MAG was calculated using CoverM in genome mode (v0.6.1, https://github.com/wwood/CoverM) by mapping clean reads from 165 metagenomes to all MAGs, with parameters “-min-read-percent-identity 0.95 -min-read-aligned-percent 0.75 -trim-min 0.10 -trim-max 0.90 -m relative_abundance”. Functional genes, including VFs, ARGs, and MGEs, were recovered and analyzed using pipelines described below, along with multiple identification tools **(Fig. S1)**.

### Prediction and localization of virulence factors

VFs were identified by searching against Virulence Factor Database (VFDB) full dataset (103) using BLASTP with an e-value cutoff of 1e-5. VFDB contains an integrated and comprehensive online resource for curated information about virulence factors of bacterial pathogens. Coding sequences (CDS) were characterized as VFs when their best hit exhibited ≥ 80% identity and ≥ 70% query coverage of the reference sequences (104). Secreted and non-secreted VFs were predicted using PathoFact (v1.0) (105), with the localization of these predicted genes identified on MGEs or chromosomes. High-confidence predictions by PathoFact (v1.0) were considered, specifically those genes annotated as VFs by Hidden Markov Model (HMM) prediction and the random forest classifier.

Functional annotation of MGEs was performed using MGEfams (https://github.com/emblab-westlake/MGEfams) with HMMER3 (106), employing the hmmsearch tool with the --cut_tc option for model-specific gathering thresholds. MGEfams comprises MGE models derived from Pfam (v 30.0) and TIGRFAMs (23), based on string matches in their functional annotations to one of the following keywords: transposase, transposon, conjugative, integrase, integron, recombinase, resolvase, conjugal, mobilization, recombination, and plasmid, as recommended previously (107).

### Identification of antibiotic resistance genes

ARGs were annotated using a range of databases. CDS were searched using HMMER3’s hmmscan with the --cut_tc option against three HMM sub-databases of ARGs: ARGfams (23), SARGfams (108), and ResFams (109), based on string matches to specific keywords. ARG-like sequences were extracted with an e-value cutoff of 1e-5. Moreover, potential ARGs were inferred using the Comprehensive Antibiotics Resistance Database (CARD) Resistance Gene Identifier (RGI) tool v6.0.3 in strict mode with the CARD database (v3.2.4) (54), as well as NCBI’s BLASTP against the Structured Antibiotic Resistance Genes (SARG) database (v3.2.1) (55) with thresholds of 30% coverage, 30% similarity, and an e-value of 1e-05. Additionally, PLM-ARG (110), based on a protein language model was employed to annotate ARGs, with ARG-like sequences having a probability greater than 0.8 identified as ARGs. Results from these approaches were integrated. Only these ARG-like sequences detected by two or more databases were identified as ARGs, and annotations only from single databases were excluded. Finally, all annotated results were consolidated and classified according to the SARG database.

The abundance of ARGs were calculated by ARGs-OAP (v3.2.4) (55), which utilizes the SARG database including 2,842 subtypes of ARGs conferring resistance to 32 classes of antibiotics. In brief, a UBLAST algorithm was employed to pre-screen ARG-like reads and 16S rRNA gene reads in metagenomic data. The ARG-like reads were then aligned against the database using BLASTX, and those with an alignment length of ≥ 25 amino acids, ≥80% similarity, and an e-value of ≤ 1e-7 were classified as ARGs.

### Distribution of high-risk antibiotic resistance genes

The risk associated with ARGs (as determined by ARGs-OAP (v3.2.4)) was assessed using ARGranker (75) based on three indicators: “human-associated-enrichment”, “gene mobility”, and “host pathogenicity”. This omics-based framework categorizes ARGs into four risk levels. ARGs that are both human-associated and mobile are classified as high-risk and further divided into “current threats” (Rank I, representing the highest risk of dissemination among pathogens) and “future threats” (Rank II, indicating a high potential for the emergence of new resistance in pathogens). ARGs that are not human-associated are assigned to Rank IV (lowest risk), while those that are human-associated but not mobile are assigned to Rank III.

### Construction of phylogenetic and structural trees

For annotated *msbA* sequences, those shorter than 400 amino acids were excluded from further analysis. The filtered sequences were clustered using CD-HIT (v4.8.1) (111) with 95% nucleotide similarity (parameters: -c 0.95 -T 0 -M 0 -G 0 -aS 0.9 -g 1 -r 1 -d 0), aligned using MAFFT (v7.505) (111), and trimmed using trimAl (v1.4.rev15) (112) with the automated1 setting. A maximum likelihood phylogenetic tree was constructed using FastTree (v2.1.11) (113) with default parameters and visualized using iTOL (v6) (114). ESMFold (115) was applied to predict the structure of each filtered *msbA* gene. A total of 202 MsbA reference protein structures were downloaded from the AlphaFold Protein Structure Database (AlphaFoldDB) (116) and the Protein Data Bank (PDB) (117). These predicted structures were aligned with those from reference proteins using Foldseek (v 9.427df8a) (118). The structural tree of MsbA was built using Foldtree (https://github.com/DessimozLab/fold_tree) (119) and visualized with iTOL (v6). The MsbA protein functions as a homodimer, and AlphaFold3 (120) was used to predict the structure of the homodimer. This predicted homodimer was aligned with the crystal structure of MsbA from *Salmonella typhimurium* (PDB ID: 3B5Z) and visualized using PyMOL (121). The complex alignment was performed using Foldseek-Multimer, calculating the TM-score normalized by both query and target length (122).

For the annotated *macA*, *macB*, and *tolC* gene sequences, which function together as a pump system, sequences from MAGs containing all three genes were selected for analysis. The *macA*, *macB*, and *tolC* sequences shorter than 300, 400, and 400 amino acids, respectively, were filtered out. Maximum likelihood phylogenetic trees of these remaining sequences were constructed in the same method as described above. A total of 141, 115, and 90 reference protein structures for MacA, MacB, and TolC, respectively, were downloaded from AlphaFoldDB and PDB. The structure of each gene was predicted using ESMFold (115) and aligned with reference proteins using Foldseek (v 9.427df8a) (118). The structural trees were built using Foldtree (119) and visualized with iTOL (v6). MacA, MacB, and TolC function as a homomeric hexamer, dimer, and trimer, respectively, forming a transmembrane pump system. The structure of this efflux pump (comprising 11 proteins) was predicted by AlphaFold3 (120), aligned with a structure from *Escherichia coli* K-12 (PDB ID: 5NIL) and visualized using PyMOL (121). The complex alignment was performed using Foldseek-Multimer (122).

### Abundance calculations of virulence factors and antibiotic resistance genes

The mapping-based mode of Salmon (v1.9.0) (123) with a “meta-flag” was used to calculate the mapping rate of the non-redundant gene catalog in each metagenome. The abundance of genes annotated as VFs were then extracted, which represented in the unit of Gene Per Million (GPM). The relative abundance of ARGs annotated by ARGs-OAP (v3.2.4), normalized to the 16S rRNA gene, was reported as gene copies per 16S rRNA gene (GP16S) (124).

Ribosomal RNAs in quality-filtered metatranscriptomic reads were removed by comparing them with rRNA sequences in the Rfam and Silva databases using SortMeRNA (v4.2.0) (125). Preprocessed reads were mapped to VFs and ARGs identified in MAGs, generating read count quantification TPM (transcripts per million) of each transcript using Salmon (v1.9.0) (123).

### Statistical analysis

Data analysis and visualization were performed using R (v4.1.3). The enrichment of different VF types and ARG types across various bacterial and archaeal phyla was calculate using Fisher’s exact test, with p-values adjusted by the Benjamini-Hochberg method. Significance was defined as an odds ratio > 1 and p < 0.05. The statistical significance of ARG abundance in cold seeps compared to other habitats, including rivers near wastewater treatment plants (water, sediment, biofilm, amphipod gut) (126) and hadal sediments (Mariana Trench, Steep Wall Site, Challenger Deep, Basin Site) (23) was assessed using the Kruskal-Wallis test. The number of VFs, ARGs, and MGEs annotated in cold seeps was compared to those in the Genomes from Earth’s Microbiomes (GEM) catalog (56) using the pairwise Wilcoxon test with Benjamini-Hochberg (BH) correction.

## Acknowledgements

This work was supported by National Science Foundation of China (No. 42376115 and No. 92351304), Natural Science Foundation of Fujian Province (No. 2023J06042), Natural Science Foundation Project of Xiamen City (No. 3502Z202373076), Scientific Research Foundation of Third Institute of Oceanography, MNR (No. 2022025 and No. 2023022) and Zhejiang Provincial High-level Talent Special Support Plan (No. 2021R51008).

Conceptualization: X.D., and T.Z. Methodology: T.Z., Y.P., and Y.H. Interpretation of data: T.Z. Visualization: T.Z. Wrote Original Draft: T.Z. Editing and review: X.D., T.Z., Y.H., Y.P., Z.D., W.S., X.X., and Y.W. All authors read and approved the final manuscript. Supervision: X.D., X.X., and Y.W. Funding Acquisition: X.D., X.X., and Y.W.

